# Quantifying the Effect of Trunk Postural Control on Reaching with Progressive Shoulder Abduction Loading in Hemiparetic Stroke

**DOI:** 10.64898/2026.09.14.750678

**Authors:** Kathleen Carolyn Suvada, Julius PA Dewald, Ana Maria Acosta

## Abstract

**Background:** Adults post hemiparetic stroke exhibit motor deficits in both the trunk and arm that impact function. In the paretic arm, involuntary coupling of torques in the fingers, wrist, and elbow during shoulder abduction, known as the flexion synergy, reduces reaching function. While there have been deficits shown to exist at the trunk such as weakness and altered coordination, it remains unclear if active trunk control compounds reaching deficits. This study examined if active trunk control with progressive shoulder abduction loading impacts reaching function in stroke and control participants.

**Methods:** Thirteen adults, 9 post hemiparetic stroke (64.11± 6.57 years) and 4 age matched controls (66.25 ± 0.96 years), completed seated reaching tasks with the trunk restrained and unrestrained while reaching with their arm coupled to the ACT-3D robotic device on a frictionless table, lifting against 25% MVT, and 50% MVT. A linear mixed effects model was used with limb as a random factor, and group, trunk restraint, and shoulder abduction load as fixed factors for quantifying changes in reaching distance expressed percent limb length (% LL).

**Findings:** Across limbs (paretic, non-paretic, control), when the trunk was un-restrained, reaching distance was reduced (*p<.05)*. Further reduction in reaching distance was observed in the paretic compared to the non-paretic and control (*p<* .*05*) limbs. Reaching velocity was fastest for controls (3.26 ± 1.52 m/s) compared to non-paretic (3.24 ± 1.19 m/s) and paretic (1.59 ± 1.32 m/s) limbs in stroke. Shoulder abduction loading had an effect on reaching distance in the paretic limb (*p<.05*) Table: .83 ± .13 % LL, 25% MVT: .77 ± .14 % LL, 50% MVT: .76 ± .15 %LL). Greater shoulder excursion occurred during non-paretic limb reaching compared to the paretic limb with an increase of 14.80 mm (*p<.05)*.

**Conclusion:** Active trunk control impacted reaching function and further reduced reaching distance in chronic hemiparetic stroke participants. However, the primary driver of reduced reaching distance was the flexion synergy. While reaching speed increased the demand at the trunk, excursions were comparable during paretic and non-paretic limb reaching, suggesting that overall trunk function was intact during this paradigm.

## Introduction

Each year, 800,000 individuals are impacted by a stroke, the primary cause of long-term disability in the United States [1]. Damage to descending corticospinal pathways and an increased reliance on cortico-reticulospinal pathways results in movement impairments in the trunk and upper limb mainly characterized by reduced reaching ability post hemiparetic stroke [2-7].

The loss of corticospinal drive and the shift to cortico-reticulospinal pathways gives rise to major impairments in the paretic limb. Weakness [10] and abnormal coupling of flexion torques in the elbow, wrist, and fingers during shoulder abduction, known as the flexion synergy [11], and hyperactive stretch reflexes [8, 9], all come at a detriment to motor function of the upper extremity. The flexion synergy, expressed as abnormal coactivation of the elbow, wrist and hand flexor muscles when supporting the limb against gravity or with added shoulder abduction loads, results in reductions in reaching distance [2], velocity [4] and work area [3], which are all further exacerbated by extensor muscle weakness. Previous work has shown that the flexion synergy is the dominant impairment impacting activities of daily living [4].The stretch reflex is believed to become hyperactive due to an increase in monoamines in the spinal cord and resulting upregulation of spinal motor neuron excitability [8, 9]. Work in the laboratory suggests that this upregulation further amplifies the drive from the cortico-reticulospinal pathways, exacerbating the flexion synergy impairment.

The vast majority of the work studying flexion synergy and reaching ability has been conducted with the trunk immobilized and affixed to a chair [3, 4, 12]. However, active trunk control, as required in activities of daily living, may exacerbate upper extremity motor control deficits and further impact reaching ability. Post stroke, studies have shown decreased elbow extension accompanied by increased shoulder and trunk movement when the trunk was actively engaged [6, 13, 14]. Interestingly, work by Khanafer et al. showed in controls that there was reduced elbow extension during reaching when the trunk was free to move compared to when trunk movement was constrained in both young and elderly participants [15]. Changes in trunk motor control during reaching after a stroke have also been documented, with trunk movement onset occurring earlier in the reach, even to targets close to the body, compared to controls where the trunk begins to move near 90% of limb length [16-21]. Note that in most of these studies, participants were not given explicit instruction to maintain trunk posture and focused instead on the reach itself, allowing varying strategies that can be incorrectly interpreted as trunk motor impairments [7, 18, 21]. Furthermore, some of these findings are confounded by unconstrained hip movement, thus preventing a clear distinction between trunk and hip motor control [15].

Studies focused on measuring in trunk motor control deficits are limited but nonetheless are important to consider as they can also impact reaching ability. Anticipatory postural adjustments (APAs) at the trunk, which are essential to the maintenance of posture [23], are delayed and reduced post-stroke [24, 25]. Furthermore, work examining multidirectional thoracic strength during maximal torque generation reveals overall weakness in comparison to control participants [26-28], with mixed evidence for lateralized weakness. Work by Dickstein et al. showed lateralized weakness in trunk flexors in the paretic side compared to the non-paretic side. In contrast, studies showed that while shifting from a flexed to a sitting posture, trunk extensor activation was comparable between the paretic and non-paretic side [5]. Yet, Tsuji et al captured reduced back extensor muscle cross-sectional area on the ipsilesional side post stroke [29]. Finally, altered trunk kinetics were found by Perlmutter et al. during maximal isometric contraction where stroke participants coupled trunk flexion with rotation and lateral flexion toward the non-paretic side as well as back extension with rotation toward the paretic side [30]. Taken together, these studies exemplify how the trunk has inherent changes in motor coordination that could impact reaching post stroke.

This study focused for the first time on the simultaneous coordination of the trunk and arm with progressive shoulder abduction loading post hemiparetic stroke. Our methodology addresses the question of whether active trunk control further compounds reaching deficits post stroke. In this novel paradigm, we tested reaching ability in stroke participants’ paretic and non-paretic arms and control participants’ dominant arm. We hypothesized that reaching ability in the paretic limb would be further reduced when simultaneously controlling trunk posture during reaching tasks on a table and while reaching with an added load in shoulder abduction analogous to the weight of the limb or heavier. Our results showed that when the trunk was free to move, there was a reduction in reaching distance for all three participant groups; however, shoulder abduction loading was the dominant factor impacting reaching distance. Results from this study show for the first time the role of the trunk in reaching ability post stroke, while confirming previous findings documenting the detrimental effects of the flexion synergy on reaching against gravity.

## Methods

Nine chronic, hemiparetic stroke participants (64.11± 6.57 years) were recruited from the Clinical Research Registry managed by the Shirley Ryan Abilitylab and Northwestern University Department of Physical Therapy and Human Movement Sciences. In addition, four age-matched controls were recruited from the community. Primary selection criteria for stroke participants included paresis confined to one side, absence of major musculoskeletal impairments in the nonparetic limb, the ability to elevate the paretic arm, and the ability to maintain an upright seated posture for prolonged periods of time. Control participants were age-matched (66.25 ± 0.96 years) and were able to maintain an upright seated posture for extended periods of time. Stroke participant clinical scores and side of paresis are listed in Table 1. Participants provided informed consent prior to participation in the study which was approved by Northwestern University’s Institutional Review Board (IRB).

**Table 1.** Stroke participant demographics including clinical scale scores: Fugl Meyer Assessment - Upper Extremity Scale (FMA-UE), Reaching Performance Scale (RPS), and Trunk Impairment Scale (TIS). Higher numbers, indicate higher performance for all scales.

| Participant | Gender (M/F) | Age (Years) | Paresis (L/R) | FMA-UE | RPS | TIS |
| --- | --- | --- | --- | --- | --- | --- |
| 1 | M | 55 | R | 20 | 12 | 12 |
| 2 | M | 67 | L | 17 | 4 | 16 |
| 3 | M | 64 | L | 42 | 32 | 15 |
| 4 | M | 62 | R | 19 | 13 | 16 |
| 5 | M | 75 | R | 9 | 13 | 16 |
| 6 | M | 54 | R | 36 | 24 | 20 |
| 7 | M | 65 | L | 23 | 15 | 15 |
| 8 | M | 69 | R | 21 | 21 | 15 |
| 9 | M | 66 | L | 7 | 0 | 12 |

### Experimental Setup and Protocol

Participants were seated in a Biodex Chair on a slanted cushion to minimize pelvic tilt and isolate the hips from the thorax. The pelvis was secured with a lap belt to minimize hip movement. For the trunk restrained condition, shoulder straps were secured to the torso to minimize trunk movement, and an additional cushion was placed against the lower back to help maintain pelvis/hip posture. During the trunk unrestrained condition, a lumbar roll placed against the lower back was used to provide support. The participant’s arm was coupled to the ACT-3D robotic device (HapticMASTER™, Motekforce Link, DIH Medical group, Amsterdam, The Netherlands) via a forearm/hand orthosis, allowing for movement in virtual reaching environments. This included a virtual table and downward vertical forces equal to 25 and 50% of the participant’s maximum voluntary shoulder abduction torque that they were asked to balance when performing the reaching task.

We used a motion tracking system (Series 2 Moiré Phase Tracking System, Metria Innovations Inc., Wauwatosa, WI) with markers attached to the trunk, scapula, humerus, and forearm/hand to obtain kinematic measurements during the reaching trials. The 3D location of bony landmarks of each segment were digitized for computation of upper body kinematics offline. Stroke participants completed two experimental sessions to test both their paretic and non-paretic arms; control participants completed the experimental paradigm once with their dominant limb for a total of n=22 limbs for all outcome measures.

Visual feedback of endpoint (3^rd^ MCP) and target position was provided in real time on a computer monitor placed directly in front of the participant (Fig. 1A). Participants were instructed to reach from the home target located in front of their shoulder with the elbow flexed to 90°(green circle in Fig. 1B), to a horizontal line placed at 110% limb length (red line, Fig. 1B). Participants were instructed and verbally encouraged to reach “as far and as fast as possible” following an auditory cue. Participants completed a total of 10 trials in each trunk restraint condition (restrained and unrestrained) and shoulder abduction loading condition (table, 25, 50% MVT) for a total of 60 trials. In the trials where the trunk was unrestrained, participants were also instructed to “keep their trunk still.” Participants were given verbal feedback of trunk posture after completion of each trial based on their ability to maintain an upright trunk posture by staying within a 5 cm circle in a transverse plane at the level of the marker placed on the sternum.

**Figure 1.**
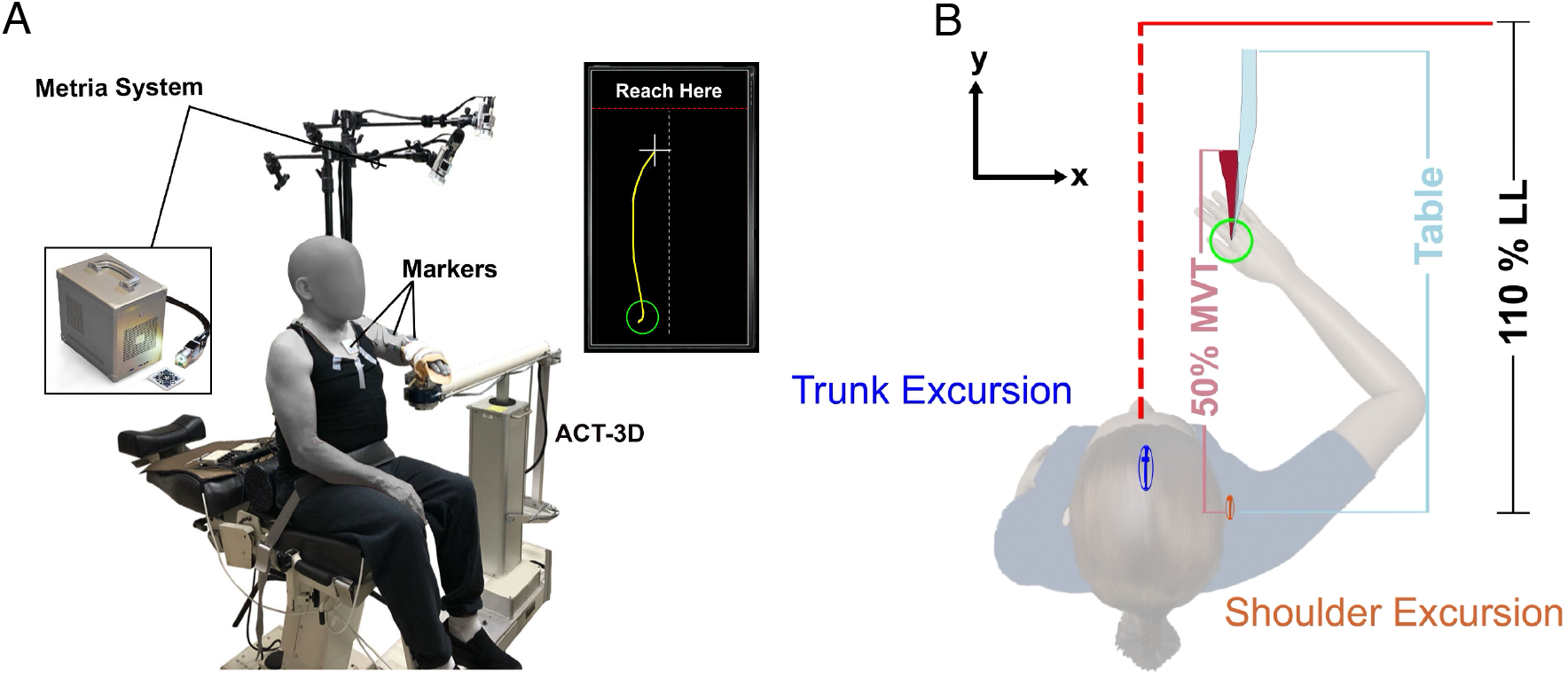
A) Experimental setup for the trunk unrestrained condition showing the ACT-3D, Metria motion capture system, and real-time visual feedback given to the participant. B) Overhead view of a representative stroke participant reaching with the paretic limb with reaching distance (RD), trunk excursion (blue), and shoulder excursion (orange). The blue shaded area depicts reaching on the table and the red shaded area red depicts lifting against 50% of MVT. The green circle represents the home target, and the red horizontal line placed at 110% LL depicts end target.

### Kinematic Data Analysis

Custom software written in Matlab (The MathWorks, Inc., Natick, MA) was used offline to obtain the 3D kinematics of all segments based on the recorded location and orientation of the Metria markers. Bone coordinate systems for the trunk, scapula, humerus, and forearm/hand were created based on the bony landmarks digitized for each segment. The 3D location of the glenohumeral joint was estimated using the a linear regression model [34, 35] based on four digitized bony landmarks of the scapula: the acromioclavicular joint, inferior angle, anterior angle of the acromion, and the medial end of the spine of the scapula [34].

Bony landmarks of the trunk (jugular notch on the sternum), shoulder (glenohumeral joint), and hand (3^rd^ metacarpophalangeal – MCP joint) were transformed to the room system as follows:

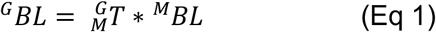

where BL is the 3×1 vector of the 3D coordinates of the bony landmark in global (G) and marker (M) coordinate frames and T is the 4x4 homogeneous transformation matrix from marker to global coordinate system.

Reaching distance was defined as the distance between the hand (3^rd^ MCP) and the glenohumeral joint in the plane of the reach. Reaching distance was normalized to limb length (%LL) to allow comparison across limbs, participants, and groups. Trunk and shoulder displacement was computed as the distance between the positions of the trunk (jugular notch) and shoulder (glenohumeral joint) at the start and end of the reach. Additional outcome measures were trunk and shoulder displacement normalized to limb length. Note that reach start timepoint was defined as the time when the endpoint velocity reached 5% of the maximum velocity prior to the first velocity peak in trials with multiple peaks, and the reach end timepoint was defined as the time when the reaching distance was maximum.

### Statistical Analysis

Differences in reaching distance (%LL), trunk excursion (cm), shoulder excursion (cm), and maximum reaching velocity were assessed using three-way mixed-effects models, with loading, restraint, and arm (control dominant, non-paretic, and paretic) as fixed factors and a limb-within-participant random intercept. A beta distribution was used for reaching distance expressed as a percentage of limb length (%LL). A Gaussian distribution was used for trunk excursion and maximum reaching velocity. Shoulder excursion was log-transformed and modeled using a Gaussian distribution. Model assumptions and goodness of fit were evaluated using Q-Q plots and simulation-based residual diagnostics. Trunk excursion had no evidence of dispersion (*p* = .354) or excess outliers (*p* = .070), although residual uniformity remained significant (*p* < .001). For shoulder excursion, residual uniformity (*p* = .053) and dispersion (*p* = .706) tests were nonsignificant, although evidence of excess outliers remained (*p* = .007). For maximum reaching velocity, there was no evidence of dispersion (*p* = .748), although residual uniformity and excess outlier tests were significant (both *p* < .001). To account for heterogeneity in residual variance, dispersion was modeled as a function of restraint, arm, and their interaction for trunk and shoulder excursion, and shoulder excursion was log-transformed to improve model fit. Remaining deviations in residual uniformity or outlier frequency indicated some departure from the assumed residual distributions; therefore, these models were retained, with inferential results interpreted considering the remaining diagnostic deviations. Post hoc comparisons of significant effects and interactions were adjusted for multiple comparisons using the Tukey method. RStudio (Posit Software, PBC) was used for all statistical analyses, with statistical significance set at *p* < .05.

## Results

### The Effect of Trunk Postural Control on Reaching

Figure 2 shows the contribution of the arm (blue), trunk (orange), and shoulder (purple) to normalized reaching distance when controls were reaching with the dominant limb (C - left panel) and stroke participants were reaching with the non-paretic (NP - middle panel) and paretic limbs (P - right column) with the trunk restrained (R) and unrestrained (U). The arm, shoulder and trunk contributions were averaged across loading conditions within limb group. Across limb groups, reaching distance was greater when the trunk was restrained (R-UR: P - 2.65% LL, NP - .96% LL, C - 1.73%LL, χ^2^(1) = 163.78, *p<* .*05)*. Notice the significant reduction in reaching distance when stroke participants reached with the paretic limb (78.7 ± 14.5%LL; right panel) compared with the non-paretic limb (95.7 ± 3.28%LL; middle panel) and controls (94.3 ± 4.09%LL; left panel).

**Figure 2.**
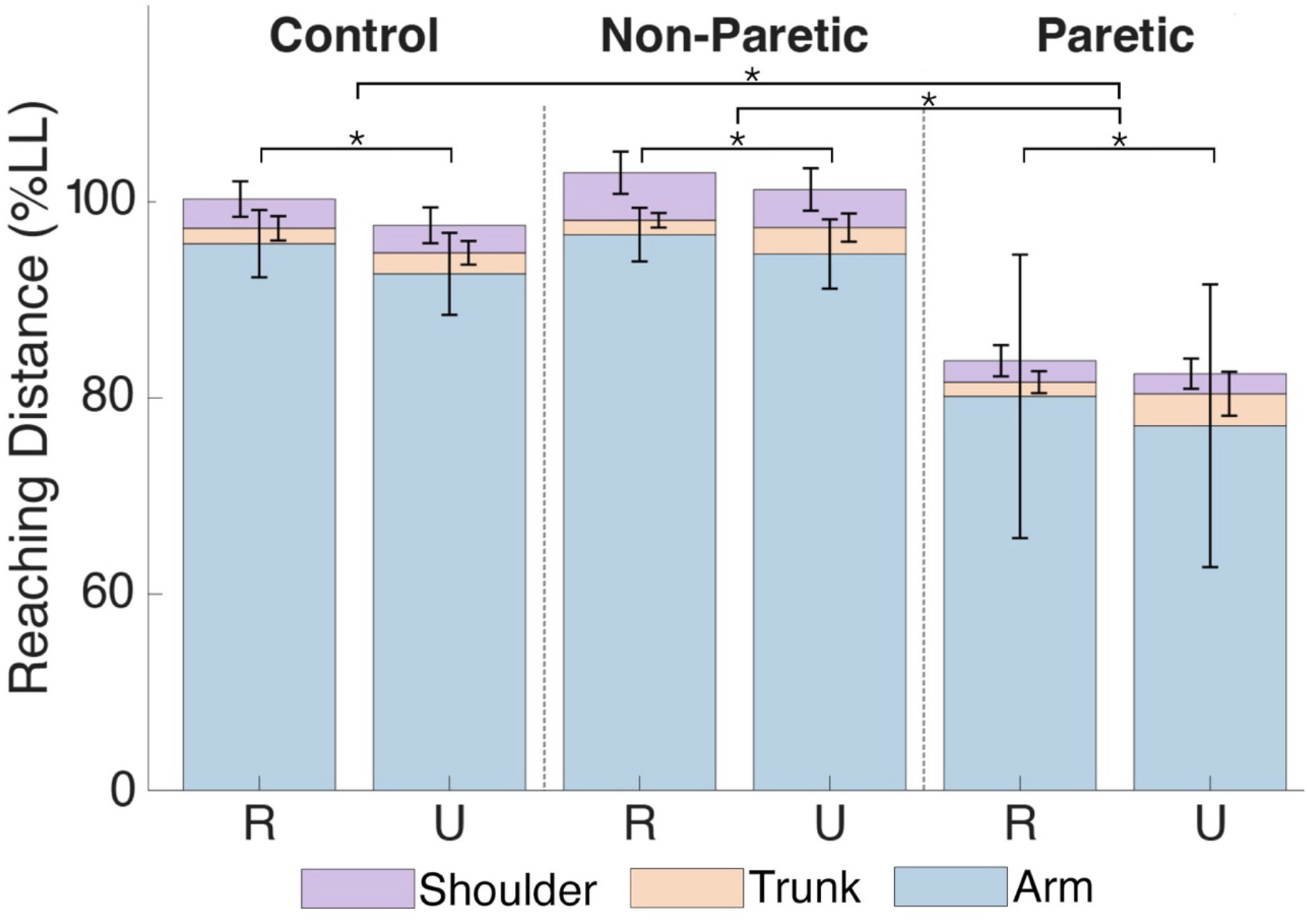
Comparison of the average contribution of the arm, trunk and shoulder across loading conditions to overall reaching distance trunk between the trunk restrained (R) and unrestrained (U) conditions; and across the paretic and non-paretic limb in stroke and dominant limb in control participants. Statistically significant (*p* <.05) reduction in reaching distance is indicated by an *.

Figure 3 depicts hand (3rd MCP joint) peak velocity during reaching. Averaged across loading and trunk restraint conditions, control participants reached with a peak velocity of 3.26 ± 1.52 m/s with the dominant limb, while stroke participants reached at 3.24 ± 1.20 m/s with the non-paretic limb and 1.59 ± 1.32 m/s with the paretic limb. Peak velocity when reaching with the paretic limb was significantly lower than in the dominant limb in controls (*p* = .015) and the non-paretic limb in stroke (*p* = .002), corresponding to approximately 49.10% and 48.48% slower reaching speeds respectively. Note that maximum velocity did not differ significantly between the dominant limb in controls and the non-paretic limb in stroke (*p* = .997).

**Figure 3.**
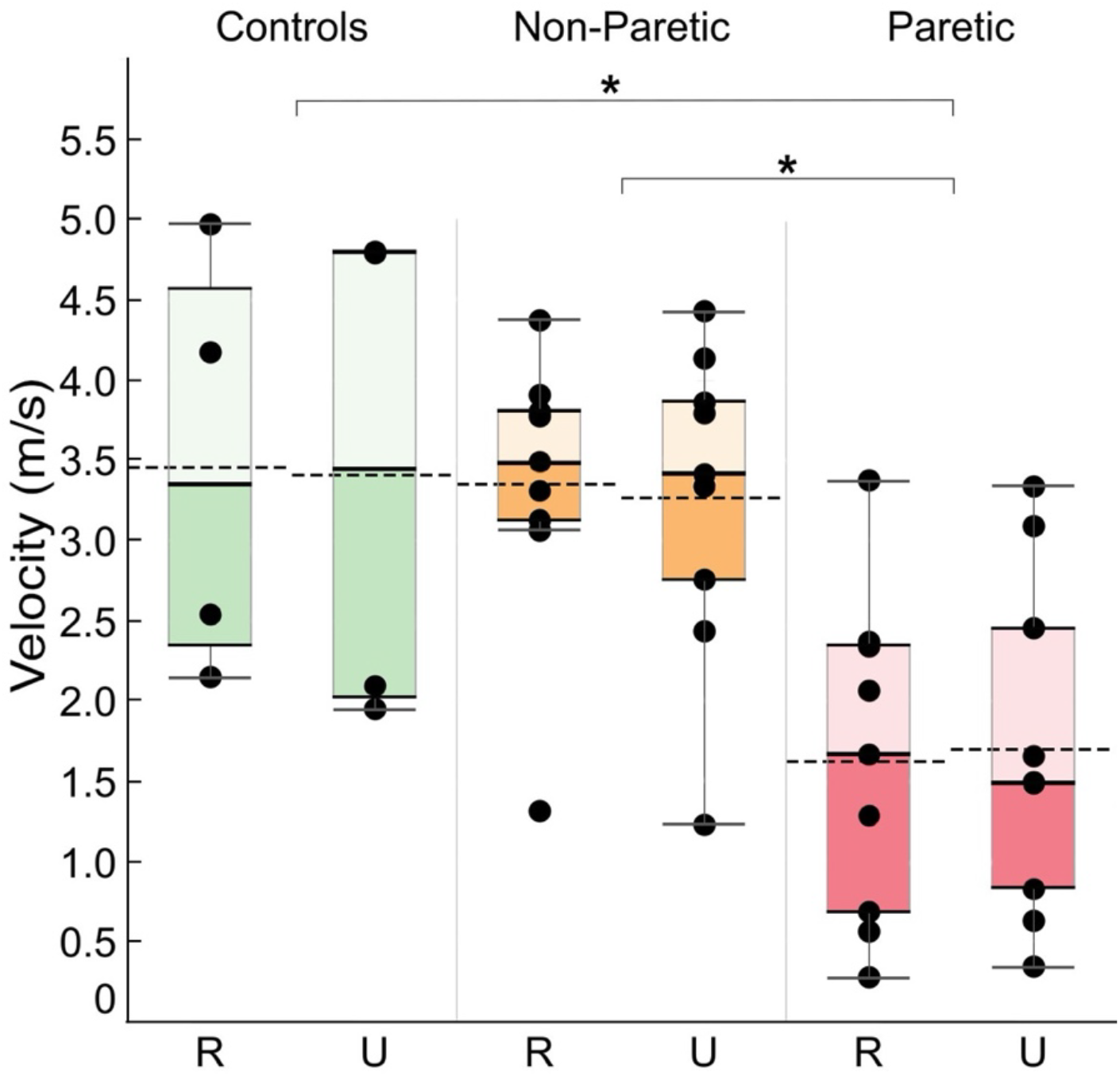
Maximum reaching velocity in m/s for controls (green), non-paretic (orange), and paretic (red) limb. R and U denote trunk restrained and unrestrained respectively. Black dashed lines represent group averages. Significance (p<.05) is indicated by an *.

Figure 4 shows reaching distance outcomes for the paretic limb and the effect of shoulder abduction load. Trunk restraint had a statistically significant effect on reaching distance across limb loading conditions (R: 85.35 ± 5.65 %LL vs UR: 82.70 ± 6.46 % LL; ES:0.821, SE: 2.15%, *p<.05*). Reaching distance was reduced by 8.1% LL with 25% MVT shoulder abduction loading and 8.3% LL with 50% MVT shoulder abduction loading in comparison to the table condition.

**Figure 4.**
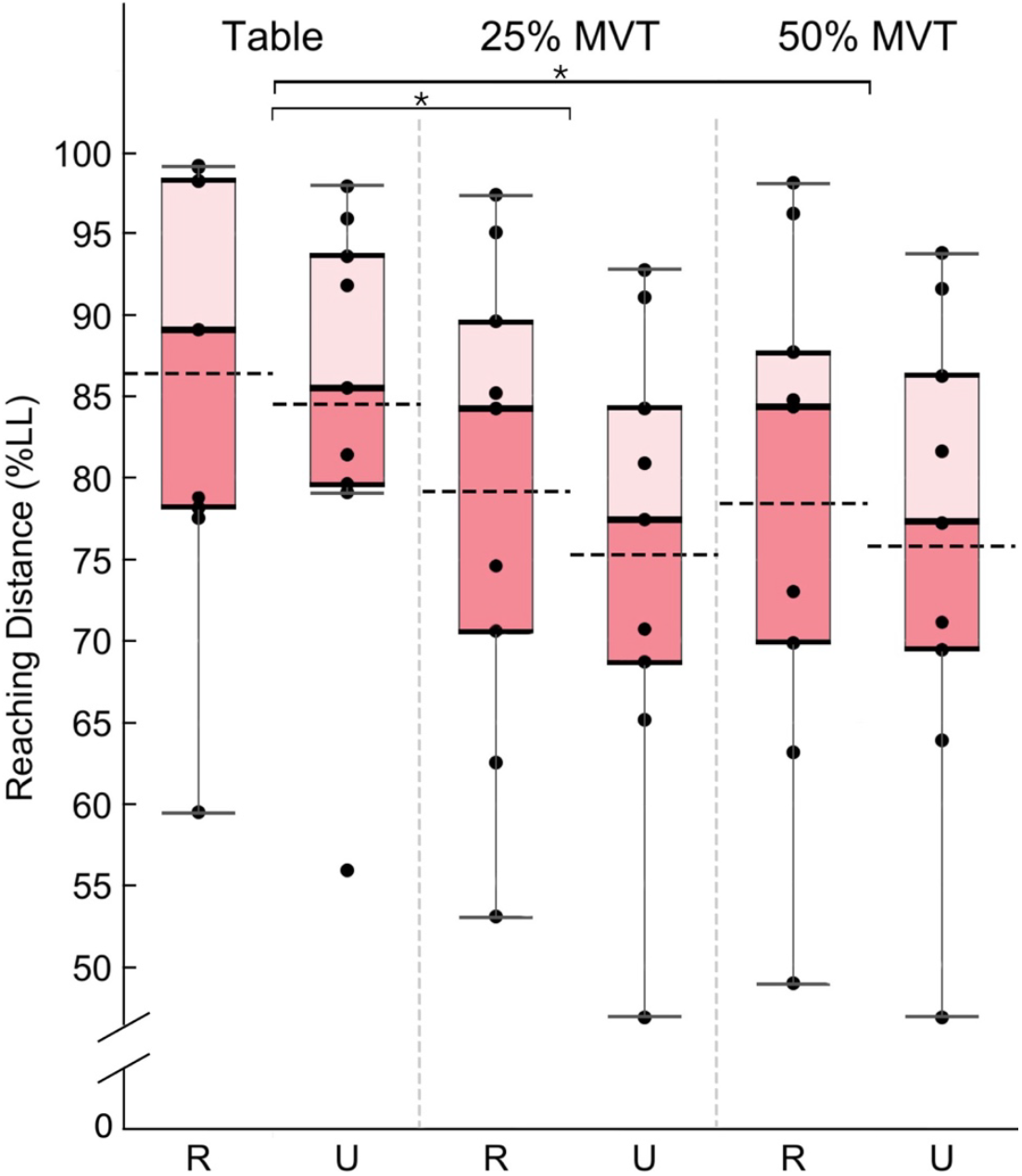
Reaching Distance normalized to limb length for all stroke participants when reaching with the paretic limb for the three loading conditions: Table, 25% MVT, and 50% MVT and with the trunk restrained (R) and unrestrained (U). Black dashed lines represent group averages. Statistically significant (*p*<.05) reductions in reaching distance are denoted by an *.

### Proximal Postural Control During Reaching

Trunk restraint had a statistically significant main effect on trunk excursion (χ^2^(1) = 6.85, *p* = .009) for all three groups, as shown in Figure 5A. Post hoc comparisons indicated that trunk excursion was significantly greater during unrestrained compared with restrained reaching for all three limb groups (all *p* < .001). Specifically, the estimated difference in trunk excursion between unrestrained and restrained reaching was 4.29 ± 0.88 cm for controls, 8.32 ± 0.70 cm for the nonparetic limb, and 11.76 ± 0.90 cm for the paretic limb, with positive values indicating greater excursion during unrestrained reaching. Figure 5B shows shoulder excursion across participant groups and trunk restraint conditions. There was an overall effect of limb group on the amount of shoulder excursion during reaching (χ^2^(2) = 9.18, *p* = .010). Post hoc comparisons indicated that shoulder excursion was greater in the restrained compared with the unrestrained condition for the NP limb, with approximately a 6.4 cm increase in shoulder excursion (*p* < .001). This difference between restraint conditions was not significant for the control (*p* = .115) or paretic (*p* = .395) limb.

**Figure 5.**
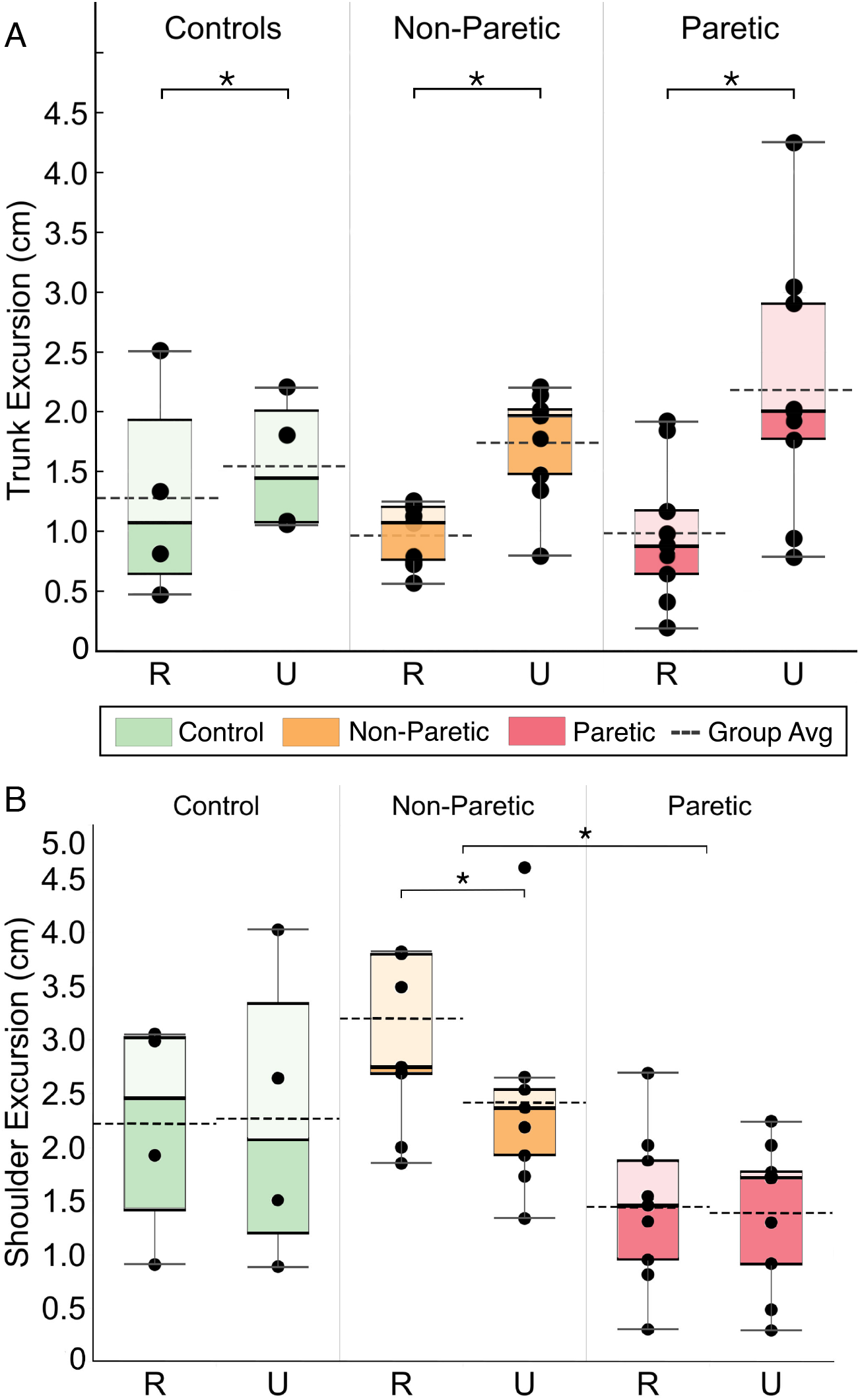
Trunk excursion (A) and shoulder excursion (B) in cm during reaching for controls (green), stroke participants: non-paretic (orange), and stroke participants: paretic (red). R and U denote trunk restrained and unrestrained respectively. Black dashed lines represent group averages. Significance (p<.05) is indicated by an *.

## Discussion

The present study evaluated the impact of active trunk control on reaching function post cortical hemiparetic stroke using a novel paradigm that required stroke participants to reach with progressive shoulder abduction loading while simultaneously controlling trunk and arm posture and arm movement. Though previous literature has characterized both a) the impact of stroke on trunk motor control and b) the impact of a stroke on reaching coordination in the presence of flexion synergy, this was the first study, to our knowledge, that examines the effect of active trunk postural control on reaching in the presence of the flexion synergy. Our overarching hypothesis was that active trunk control would further compound reaching deficits already present due to the flexion synergy. This central hypothesis was supported by our data that showed reduced reaching ability as measured by reaching distance in the active trunk posture conditions.

Interestingly, we observed reduced reaching distance across all participant groups and tested limbs when actively controlling trunk posture. For stroke participants, active control of trunk posture further compounded reaching deficits due to the flexion synergy in the paretic limb. However, the effect of trunk restraint (ES: 0.821) on reaching function was only a fraction of the effect of the flexion synergy (ES: 1.95), highlighting the devastating impact of the loss of corticospinal drive and increased reliance on cortico-reticulospinal pathways. Our findings agree with past literature that present the flexion synergy as the dominant impairment to reaching function post stroke [4].

Overall, control and stroke participants were similarly able to maintain their trunk posture during reaching with less than 5 cm trunk excursion across experimental conditions. Note that individuals with stroke allowed greater trunk excursion during reaching with the paretic limb compared to reaching with the non-paretic limb or the dominant limb in controls. When comparing normalized (to limb length) trunk and shoulder excursion during reaching, it was clear that the proximal body (trunk and shoulder) only contributed minimally to the reach even in stroke participants and therefore the task was primarily completed through shoulder and elbow movement. This is in contrast with past studies that did not give explicit instruction to maintain trunk posture while reaching to targets in front of the trunk and therefore may have underestimated their ability to control their trunk posture. Interestingly, average trunk excursion during reaching with the non-paretic and paretic limbs in stroke participants was comparable, demonstrating their ability to maintain the trunk posture even with the non-paretic limb moving at speeds nearly twice those of the paretic limb, and resulting in greater perturbations to the trunk. Trunk postural control may be, in the context of this reaching task, more intact due to multiple backup motor pathways, descending bilaterally, and the preferred reliance on brainstem versus cortical pathways for motor control [32, 33]. This may be a reason for the trunk exhibiting less impairment compared to the arm. This suggests that the greater trunk excursion while reaching with the paretic limb is a compensatory mechanism to account for reaching impairments imposed by the flexion synergy and not diminished trunk postural control as indicated in previous studies [7].

An unexpected behavior captured in our protocol was greater shoulder protraction during reaching when the trunk was restrained to the chair compared with when the trunk was free to move, with the non-paretic limb demonstrating the greatest shoulder protraction across limbs. We speculate that participants use the chair as a ground to maximize reaching velocity through the shoulder and elbow muscles. Specifically, when the trunk is restrained, participants may maximally activate not only the muscles that rotate the shoulder in horizontal flexion, but also the muscles that protract the shoulder forward in the direction of the reach [27, 29, 34].

This study quantified for the first time, the effect of stroke on trunk postural control and its interaction with the flexion synergy in a reaching task. While past studies revealed specific impairments in trunk postural control due to hemiparetic stroke, their measurements were made in isolation or without specific instruction to engage trunk postural muscles. In daily life, stroke survivors are coordinating their trunk and arm together under the influence of the flexion synergy, thus grounding the results from this study that demonstrate that active trunk control is largely preserved following stroke at least in the context of a ballistic reaching task with the paretic and non-paretic limbs, while the flexion synergy is the main contributor to deficits in reaching ability. Results from studies that either restrain the trunk or provide no instruction regarding trunk posture need to be carefully interpreted, as they may conflate limitations in reaching ability with deficits in trunk postural control or overestimate reaching ability when allowing individuals to ground their trunk against a surface as would happen when strapping the participants to the back of the chair.

This study has several limitations that can influence the results. First, while marker-based motion capture is proven to be a valid and useful method for measuring human joint kinematics, its application to the shoulder specifically to track scapular motion is less common. However, we believe that careful placement of the markers to minimize skin movement on the scapula coupled with computation of the center of the glenohumeral joint based on scapular bony landmarks resulted in representative rotation and translation of the joint. Second, the demands of the reaching task studied here may not have been sufficient to elicit trunk specific deficits. Previous studies examining trunk weakness and abnormal coupling following stroke required maximal effort, thereby characterizing the limits of trunk postural control. In the present study we were interested in characterizing trunk and arm postural control using a task that is representative of activities of daily living. Future work may expand these results by examining tasks with greater demand on trunk postural control potentially through reaching in different directions greater loads on the arm or even loading the trunk. Finally, the average level of trunk impairment in the participant pool as measured with the Trunk Impairment Scale was mild (score = 15.22/23), indicating mild trunk function. It is possible that this biased our results, and could be addressed in future studies by recruiting participants who have greater impairments in trunk function.

## Conclusion

This study was the first to examine reaching ability while combining active trunk control with progressive shoulder abduction loading post hemiparetic stroke. Our results showed that active trunk control further reduced reaching distance in both limbs of stroke participants, increased trunk excursion when un-restrained in both limbs, and increased shoulder excursion in the non-paretic limb specifically when the trunk was restrained. The findings from this work support current rehabilitation practices that integrate the trunk into the physical therapy interventions to help restore reaching function to after a stroke and provide interesting considerations when designing experiments to examine reaching ability.

## Notes

### Competing Interest Statement

The authors have declared no competing interest.

### Funding

- NIH-NICHD: R01HD039343
- NIH-NICHD: T32HD101395

